# From Prediction to Action: Dissociable Roles of Ventral Tegmental Area and Substantia Nigra Dopamine Neurons in Instrumental Reinforcement

**DOI:** 10.1101/2022.08.15.501890

**Authors:** Kurt M. Fraser, Heather J. Pribut, Patricia H. Janak, Ronald Keiflin

**Affiliations:** Department of Psychological & Brain Sciences, Krieger School of Arts and Sciences, Johns Hopkins University, Baltimore, MD 21218, USA; The Solomon H. Snyder Department of Neuroscience, Johns Hopkins School of Medicine, Johns Hopkins University, Baltimore, MD 21205, USA; Kavli Neuroscience Discovery Institute, Johns Hopkins University School of Medicine, Baltimore, MD 21205, USA; Department of Psychological & Brain Sciences, University of California, Santa Barbara, Santa Barbara, CA 93106, USA

## Abstract

Reward-seeking requires the coordination of motor programs to achieve goals. Midbrain dopamine neurons are critical for reinforcement and their activation is sufficient for learning about cues, actions, and outcomes. Here we examine in detail the mechanisms underlying the ability of ventral tegmental area (VTA) and substantia nigra (SNc) dopamine neurons to support instrumental learning. By exploiting numerous behavioral tasks in combination with time-limited optogenetic manipulations, we reveal that VTA and SNc dopamine neurons generate reinforcement through separable psychological processes. VTA dopamine neurons imbue actions and their associated cues with motivational value that allows flexible and persistent pursuit whereas SNc dopamine neurons support time-limited, precise, action-specific learning that is non-scalable and inflexible. This architecture is reminiscent of actor-critic reinforcement learning models with VTA and SNc instructing the critic and actor, respectively. Our findings indicate that heterogeneous dopamine systems support unique forms of instrumental learning that ultimately result in disparate reward-seeking strategies.

## INTRODUCTION

Midbrain dopamine neurons encode reward prediction errors, a fundamental parameter in reinforcement learning^1–3^, and their activation promotes learning about events leading to reward ^4–9^. Although early studies reported relatively broad and homogeneous responses to unexpected rewards across midbrain dopamine cell groups — including the ventral tegmental area (VTA) and the substantia nigra (SNc)—^10–12^, more recent studies have demonstrated considerable heterogeneity in the response pattern of different dopamine subsystems, particularly in relation to non-reward variables^13–20^. For instance, dopamine neurons in the medial VTA signal cue and outcome related information, whereas dopamine neurons located in the lateral VTA and SNc appear to encode motor parameters in reward-guided tasks^16,17^. Moreover, these different dopamine cell groups (VTA and SNc) have largely dissociable projection targets, with VTA dopamine (VTA^DA^) neurons projecting predominantly to the limbic ventromedial striatum, and SNc dopamine (SNc^DA^) neurons projecting predominantly to the sensorimotor dorsolateral striatum^21–23^. The dissociable response profiles and the relatively segregated anatomical targets of VTA and SNc dopamine neurons strongly suggest a functional dissociation between these neuronal populations. In line with this idea, cues paired with optogenetic activation of either VTA or SNc dopamine neuron acquire qualitatively different motivational properties (selective approach vs. general locomotion respectively)^23^.

Importantly, this regional specialization of dopamine neurons was evident only in Pavlovian conditioning preparations, in which subjects can anticipate —but not control— the delivery of optogenetic dopamine stimulation. This contrasts with the seemingly uniform role for dopamine neurons in self-stimulation preparations, in which the activation of dopamine neurons is contingent on an instrumental response. Indeed, rats will avidly press a lever if this results in the activation of their dopamine neurons, regardless of whether stimulation is delivered to the VTA or SNc^4,8,23,24^.

Why is there strong evidence for functional specialization of VTA and SNc dopamine neurons in Pavlovian but not instrumental reinforcement learning? Although it may be the case that VTA and SNc dopamine neurons contribute uniformly and undistinguishably to instrumental reinforcement, an alternative and more likely hypothesis is that this functional homology is only apparent in reduced or constrained scenarios. Although activation of VTA or SNc dopamine neurons favors the repetition of an instrumental response, the underlying motivational processes engaged by these two neural populations might differ. The purpose of this study was to test this hypothesis of a functional heterogeneity of VTA and SNc dopamine neurons in instrumental reinforcement. For this purpose, rats were trained to lever-press for optogenetic stimulation of VTA or SNc dopamine neurons; these rats were then subjected to different behavioral assays and manipulations designed to probe the nature and content of the processes governing their instrumental response.

## RESULTS

To achieve selective control of midbrain dopamine neurons, we injected a Cre-dependent viral vector for the expression of channelrhodopsin (ChR2) into the VTA or SNc of transgenic TH-Cre rats and implanted an optic fiber aimed at those regions (**Figure 1**). All behavioral procedures were conducted 3 - 6 weeks after surgeries. In all experiments described below, rats were initially trained to press one of two levers (designated as active) to obtain a brief optogenetic activation of VTA or SNc dopamine neurons (2s stimulation, at 20Hz). Rats rapidly acquired reliable and stable self-stimulation responding (within 3-11 sessions; data not shown). Responding on the inactive lever was low for both groups (VTA: 0.844 ± 0.236; SNc: 1.219 ± 0.312 responses per hour [mean ± s.e.m.] after initial ICSS acquisition; U = 235.5, P = 0.276; 95% CI [-0.342, 1.135]; effect size = 0.182) and remained negligible throughout the different manipulations. Therefore, for simplicity, we will only present active-lever presses.

**FIGURE 1:**
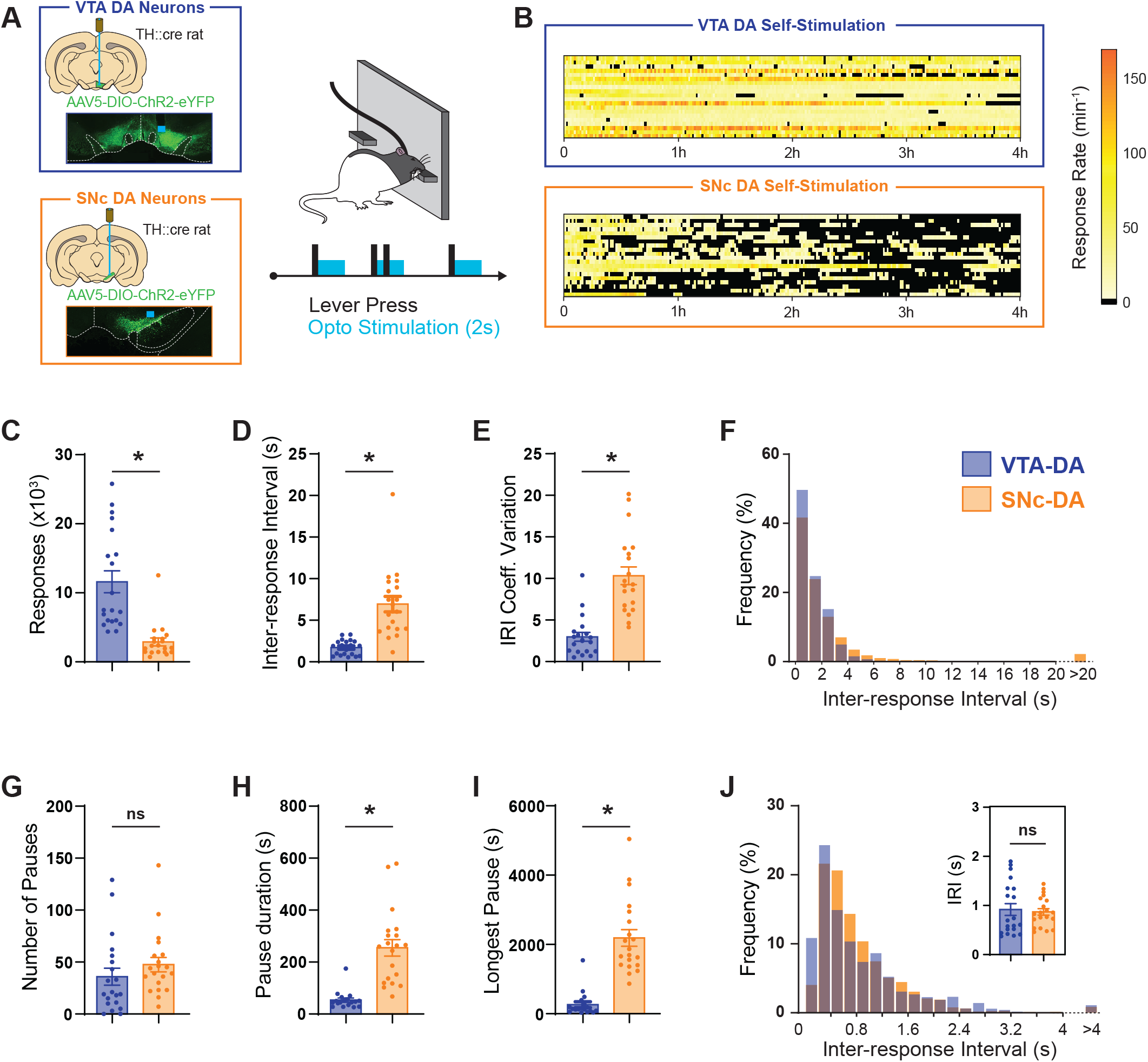
Self-stimulation of VTA or SNc dopamine neurons produces different patterns of operant responses. **A**. TH::Cre rats were made to express ChR2 in either VTA (n=20) or SNc (n=20) dopamine neurons. Responding on the active lever resulted in a 2s optoactivation of the targeted dopamine neurons. **B**. Heat maps of the rate of operant responding throughout the 4h sessions, in animals self-stimulating VTA-or SNc-DA neurons. Each line represents a different animal. **C**. Total responses on the active lever. **D**. Average inter-response interval (IRI) **E**. Coefficient of variation of IRI. **F**. Frequency distribution of IRIs (as percentage of total responses). G. Number of within-session pauses in operant responding (pause defined as a IRI>20s). **H**. Average pause duration **I**. Longest pause in a session. **J**. Frequency distribution of IRIs during a 4-min peak responding period. (* P<0.001 Mann-Whitney U-tests). Error bars = SEM.

### Self-stimulation of VTA or SNc dopamine neurons generates different patterns of instrumental responding

Response-contingent activation of VTA or SNc dopamine neurons resulted in striking differences in instrumental response patterns (**Fig.1**). Subjects self-stimulating VTA^DA^ neurons displayed high rates of responding throughout the 4h session. In contrast, responding for SNc^DA^ neurons activation was characterized by bouts of vigorous responding interrupted by long pauses. This resulted in significant group differences in the total number of operant responses (U = 16, P < 0.001; 95% CI [5988.18, 12595.94]; effect size = 0.920) and the average inter-response intervals (IRI; U = 16.00, P < 0.001; 95% CI [3.78, 6.92]; effect size = 0.920). To assess the regularity of responding within a session, we calculated for each subject the coefficient of variation of IRIs. Subjects self-stimulating SNc^DA^ neurons displayed higher coefficient of variation (U = 21.00, P < 0.001; 95% CI [4.87, 9.44]; effect size = 0.895), consistent with the irregular pattern of responding in this group. Despite significant differences between VTA and SNc groups in average IRIs, the relative distribution of IRIs appears remarkably similar in both groups, with the vast majority (>90%) of responses occurring in rapid succession (≤ 4s IRI). Where VTA and SNc groups differ most profoundly is in the duration of their instrumental pauses (pauses were arbitrarily defined as a period of 20s or longer that separates two instrumental responses). These pauses were equally frequent in both groups (T = 0.698, P = 0.490; 95% CI [-13.79, 28.37]; effect size = 0.335), but lasted much longer in subjects self-stimulating SNc^DA^ neurons (U = 9.00, P < 0.001; 95% CI [144.36, 264.51]; effect size = 0.955). Analysis focused on periods of peak responding in a session confirmed that when actively engaged in responding both VTA and SNc groups responded at the same rate (U = 183.00, P = 0.655; 95% CI [-0.35, 0.23]; effect size = 0.085).

To determine whether these different response patterns could reflect different reward intensities induced by VTA vs. SNc dopamine neuron stimulation, we systematically reduced the intensity of the stimulation by reducing the laser power in a subset of VTA^DA^ self-stimulating rats. This manipulation failed to reproduce the response pattern that characterizes SNc^DA^ neurons self-stimulation (**Fig.S1**). This suggests that, rather than producing different intensities of reinforcement, activation of VTA^DA^ or SNc^DA^ neurons engages different reinforcement processes.

### Imposed timeouts abolish instrumental responding for SNc, but not VTA dopamine neurons self-stimulation

The irregular pattern of responding observed in the SNc group suggests that, unlike subjects in the VTA^DA^ group, animals self-stimulating SNc^DA^ neurons might lack the motivation to approach the active lever and initiate bouts of responding (this is despite the fact that both groups respond avidly for dopamine self-stimulation once a bout has been initiated). However, given the established role of the nigrostriatal dopamine pathway in motor functions^17,25,26^, an alternative explanation for the irregular and interrupted pattern of responding in the SNc^DA^ group is that the activation of SNc^DA^ neurons produces motor effects that are incompatible with sustained high rates of instrumental responding^27^. The following experiment was intended to tease apart the contribution of these two potential mechanisms (motivation to initiate responding vs. competing motor effects).

After initial self-stimulation training (as described above), rats were tested in a session in which each press on the active lever resulted in the optogenetic activation of VTA (n = 15) or SNc (n = 16) dopamine neurons, and was immediately followed by the retraction of both active and inactive levers for a duration of 12s. After this imposed timeout period, both levers were again extended and rats could press the active lever for a subsequent stimulation (**Fig. 2**). This reinforcement schedule limits the number of stimulations that can be obtained in a session; it also allows for potential stimulation-induced motor effects to dissipate before the next opportunity to respond. We reasoned that if motor effects are responsible for the reduced responding in the SNc group, then the imposed timeouts should mitigate these motors effects and tend to equalize responding between the two groups. On the other hand, if reduced responding in the SNc group is due to a reduced motivation to initiate responding after a pause, then the imposed timeouts should further reduce responding in that group, as every lever press now requires subjects to initiate responding by (re)approaching the lever (i.e. continuous high frequency presses are no longer possible).

**FIGURE 2:**
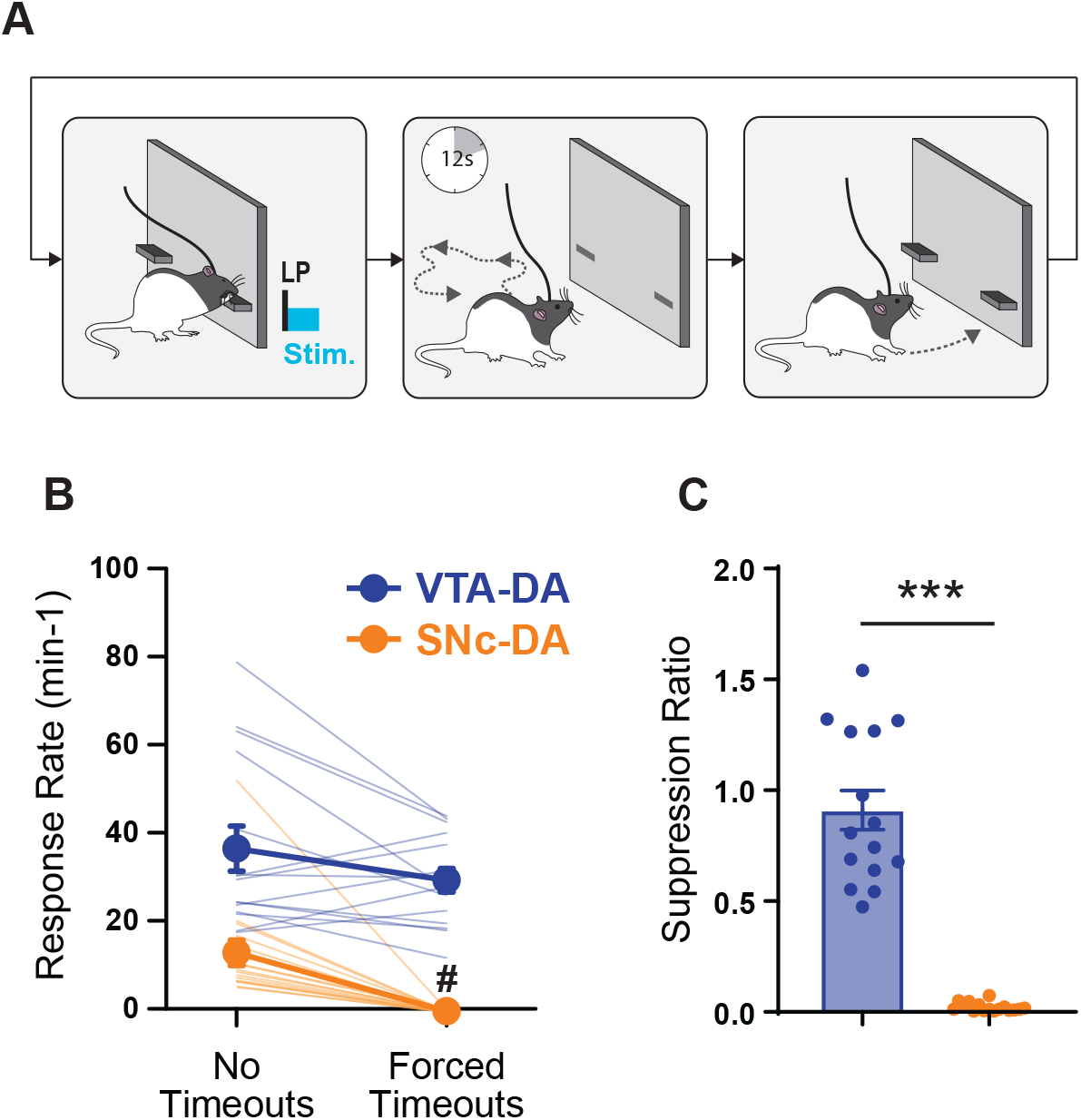
Imposed timeouts abolish responding for SNc^DA^ but not VTA^DA^ neuron stimulation. **A**. Behavioral paradigm. Each response on the active lever results in the optical stimulation of VTA^DA^ (n=15) or SNc^DA^ (n=16) neurons, and triggers the retraction of both levers for a period of 12s (imposed timeout). **B**. Response rate in absence or in presence of imposed timeouts. **C**. Suppression ratio induced by imposed timeouts. (# P< 0.001 Wilcoxon Z-test [No Timeouts vs Imposed Timeouts]; * P<0.001 Mann-Whitney U-test [VTA^DA^ vs. SNc^DA^]). Error bars = SEM.

To facilitate comparison between continuous reinforcement and imposed timeouts sessions, we calculated the response rate during the period when the levers were present (when a lever press response could actually be produced). The imposed timeouts significantly decreased response rate in animals self-stimulating SNc dopamine neurons (Z = -3.516, P < 0.001; 95%CI [8.890; 20.167; effect size = 1.00), but not in animals self-stimulating VTA dopamine neurons (Z = -1.477; P = 0.151; 95% CI [-0.980, 15.367]; effect size = 0.333). To directly compare the two groups, we calculated for each subject a suppression ratio, defined here simply as the response rate during the imposed timeout session divided by the response rate during continuous reinforcement. We found that compared to animals self-stimulating VTA dopamine neurons, animals self-stimulating SNc dopamine neurons were more sensitive to imposed timeouts (U = 0.000; P < 0.001; 95% CI [0.726, 1.060)]; effect size = 1.00). This strongly suggest that, compared to animals in the VTA group, animals self-stimulating SNc dopamine neurons express reduced motivation to reengage with the lever and reinitiate responding after a pause.

### Topological changes in response requirement abolish instrumental responding for SNc, but not VTA dopamine neurons self-stimulation

Our result thus far suggest that self-stimulation of dopamine neurons in the VTA or SNc engages qualitatively different instrumental processes. While subjects in both groups are capable of highly stereotyped responding within a bout, SNc subjects appear deficient in their ability to return to the lever and reinitiate responding after a pause. An important distinction between within-bout responding and bout initiation is that only the later requires a flexible approach strategy, since for every bout initiation (and depending on animals’ initial location and position in the chamber) a new set of actions is required to approach and reach the lever^28^. In the following experiment, we decided to further interrogate the flexible approach component of instrumental responding by imposing environmental constraints that required subjects to perform a new -topologically different-response to reach and press the lever.

Rats initially trained to press a lever to self-stimulate VTA (n = 10) or SNc (n = 8) dopamine neurons, were tested in two probe sessions: in one session the lever remained in its standard position, in another session the lever was raised by 8 cm (order counterbalanced, 3 days of retraining between probe sessions). The raised lever was still within reach but activating the lever now required a different set of motor commands, resulting in a different body posture (crouching in the standard lever position vs. rearing in the elevated lever position, **Fig. 3**). Importantly, optogenetic stimulation was not delivered in these probe tests in order to avoid the confounding effects of within-session reinforcement of the new response. Throughout this experiment, only the active lever was presented in order to prevent the potential transfer of responding between levers. Moreover, to facilitate the detection of the lever, a stimulus light located above the lever was continuously illuminated (during training and probe test). Performance on this probe session was compared with the standard probe session in which the lever was in its usual position. A 2-way mixed ANOVA (Group x Session) found no main Group effect (F_1,16_ = 4.093, p = 0.060) but a significant Session effect (F_1,16_ = 32.029, p < 0.001) and Group x Session interaction (F_1,16_ = 4.971, p = 0.04). Planned post hoc comparisons indicated that while SNc and VTA group did not differ in a standard extinction session (T = 0.179; p = 0.859; 95% CI [-23.00, 19.10]; effect size = 0.075), the relocation of lever induced a reduction in responding that was much more pronounced in the SNc group, resulting in a significant difference between these groups (T = 2.974; p = 0.006; 95% CI [-46.726, -15.146]; effect size = 0.824). This indicates that, unlike subjects in the SNc group, subjects in the VTA group were able to improvise a new set of actions to reach and activate the relocated lever.

**FIGURE 3:**
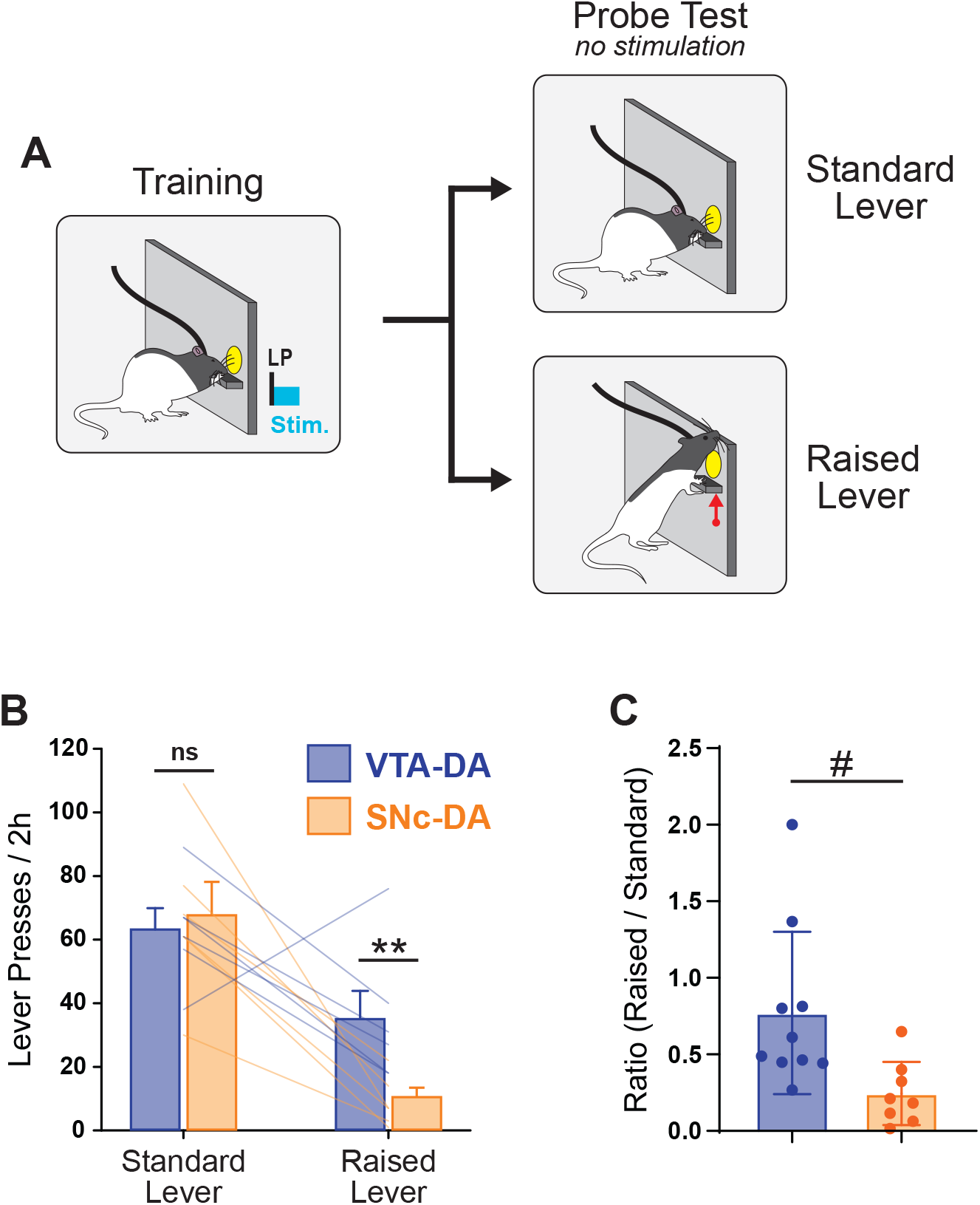
Lever relocation, and modified response requirement, has moderate effects on instrumental responding for VTA^DA^ stimulation, but strongly disrupts instrumental responding for SNc^DA^ stimulation. **A**. Rats trained to self-stimulate VTA (n=10) or SNc (n=8) dopamine neurons were tested in two nonreinforced probe sessions. For one probe session, the position of the active lever was raised, thereby imposing a new set of motor commands to reach and activate the lever. **B**. Instrumental responses during probe sessions, with standard or relocated (raised) lever position. **C**. Suppression ratio induced by lever relocation. (*P<0.01 T-test; # P<0.01 Mann-Whitney U-test, 95% CI [-0.897, -0.213], effect size = 0.8.). Error bars = SEM.

### Cues paired with VTA but not SNc dopamine neurons stimulation increase responding in progressive ratio

Our results thus far indicate that, compared to animals in the SNc group, animals in the VTA group express a higher propensity to approach the active lever operandum. This suggests that in this last group, the circumstances (spatial location and/ or environmental stimuli) surrounding optogenetic stimulation have acquired some incentive properties that compel subjects to approach the active lever and initiate responding. In addition to their ability to motivate approach, another defining property of incentive stimuli is that animals will work to obtain those stimuli, even in the absence of the primary reward they signal. Therefore, in this next experiment, we decided to formally test the incentive properties acquired by phasic stimuli paired with VTA or SNc dopamine neurons self-stimulation, by testing their ability to maintain instrumental responding in the (near) absence of actual optogenetic stimulation.

Rats were trained to self-stimulate VTA (n = 6) or SNc (n = 6) dopamine neurons, and each stimulation coincided with the presentation of a brief audiovisual cue (white noise and chamber illumination; 0.4s; **Fig. 4**). To facilitate the association between this cue and the optogenetic stimulation of dopamine neurons, we progressively increased the response requirement (from FR1 to RR2), which reduced the response-stimulation contingency, while maintaining a maximal cue-stimulation contingency—effectively increasing the relevance of the cue. Moreover, to ensure equal number of cue-stimulation pairings in both groups, we imposed a maximal number of stimulations per session, which all animals completed (**Fig. S2**).

**FIGURE 4:**
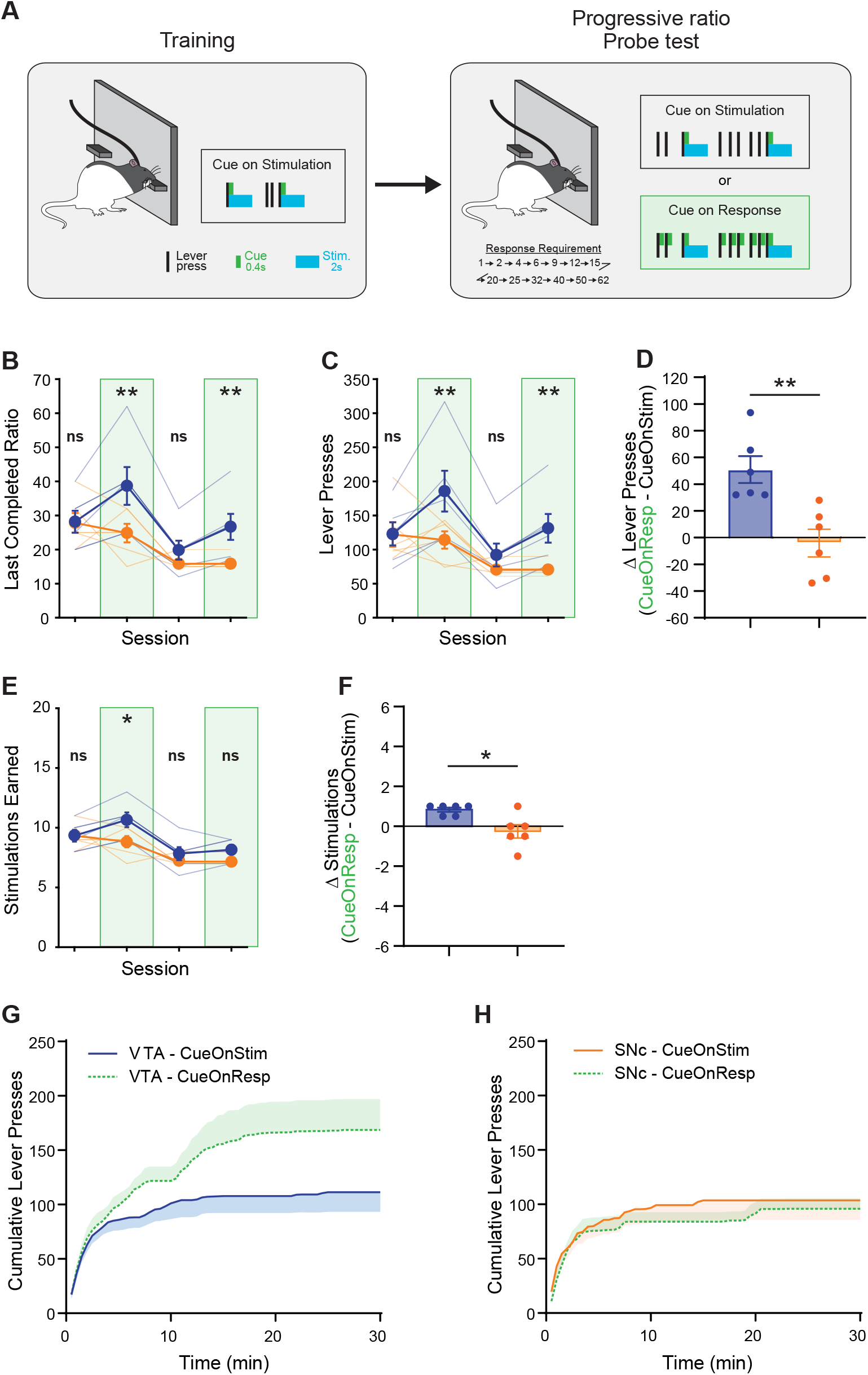
Cues paired with VTA^DA^ but not SNc^DA^ neuron stimulation increases responding in progressive ratio. **A**. Behavioral paradigm. During VTA^DA^ (n=6) or SNc^DA^ (n=6) self-stimulation training, every optostimulation was paired with a brief audiovisual cue. Responding was then tested in progressive ratio sessions, in which the cue continued to accompany every stimulation (cue on stimulation; session 1 and 3), or was contingent on every active lever press (cue on response; session 2 and 4). **B, C** Highest completed ratio **(B)**, and total lever presses **(C)** in progressive ratio probe sessions. **D**. Difference score, obtained by subtracting the average responding in sessions 1 and 3 from the average responding in sessions 2 and 4. **E-F**. Number of stimulations earned **(E)** and difference score **(F). G-H**. Response time course (cumulative responses) during progressive ratio probe tests for VTA **(G)** and SNc **(H)** groups. (*P<0.05, **P<0.01 T-tests [VTA vs SNc]). Error bars = SEM.

Finally, animals were tested in four behavioral sessions in which optogenetic stimulation was delivered according to a progressive ratio (PR) schedule—a situation in which the vast majority of responses do not result in optogenetic stimulation. In two of those PR test sessions (session 1 and 3), the audiovisual cue continued to be presented on every stimulation (as in training). In the other two PR test sessions (session 2 and 4), the audiovisual cue was contingent on every response on the active lever. A 2-way mixed ANOVA (Group x Session) conducted on the number of responses on the active lever found no main Group effect (F_1,10_ = 3.161, P = 0.106), but a significant Session effect (F_3,30_ = 16.937, P < 0.001) and Group x Session interaction (F_3,30_ = 5.284, P = 0.013; Greenhouse-Geisser corrected). Post-hoc tests revealed that VTA and SNc group did not differ in sessions in which the cue was presented only during receipt of optogenetic stimulation (Ps > 0.398). However, the introduction of response-contingent cue increased responding only for the VTA group (S1 vs. S2 and S3 vs. S4: Ps < 0.010) and not for the SNc group (Ps > 0.591), resulting in significant group differences on those session (Ps < 0.030). This increase in responding observed in the VTA group resulted in a very modest but significant increase in the number of stimulations obtained by that group (**Fig. 4 E-F**). The fact that only the VTA group benefited from phasic, response-contingent, optogenetic stimulation-paired cues, indicates that cues paired with the activation of VTA, but not SNc dopamine neurons acquired incentive properties.

### Activation of VTA but not SNc dopamine neurons sustains heterogeneous instrumental sequences

Our results thus far indicate that while the activation of either VTA or SNc dopamine neurons serves as a potent reinforcer of instrumental actions, only in the VTA group does the “state” (location and/or environmental stimuli) associated with dopamine stimulation acquire some incentive value. An evolutionary advantageous property of a state-value function is that, by signaling when the prospect of reward has increased, it can guide animals through the acquisition of complex instrumental sequences leading to a primary reinforcer^29,30^. In this experiment, we tested to which extent subjects self-stimulating VTA or SNc dopamine neurons could acquire an increasingly complex action sequence to obtain optogenetic stimulation.

Rats were initially trained to press a lever to obtain optogenetic stimulation of either VTA (n=13) or SNc (n=14) dopamine neurons. After five or six sessions (post initial acquisition) under this reinforcement schedule, rats were then required to perform a sequence of two instrumental actions to obtain an optogenetic stimulation. Specifically, they were required to press one lever (the previously inactive lever) to gain access to another lever (the previously active lever) that they could then press to obtain the stimulation (i.e. a seeking–taking reinforcement schedule). After three sessions under this reinforcement schedule, the required instrumental sequence was further extended. To obtain an optogenetic stimulation, rats had to perform a sequence of three instrumental actions, starting with a nosepoke, then a press on a first (seeking) lever, and finally a press on a second (taking) lever (**Fig. 5**).

**FIGURE 5:**
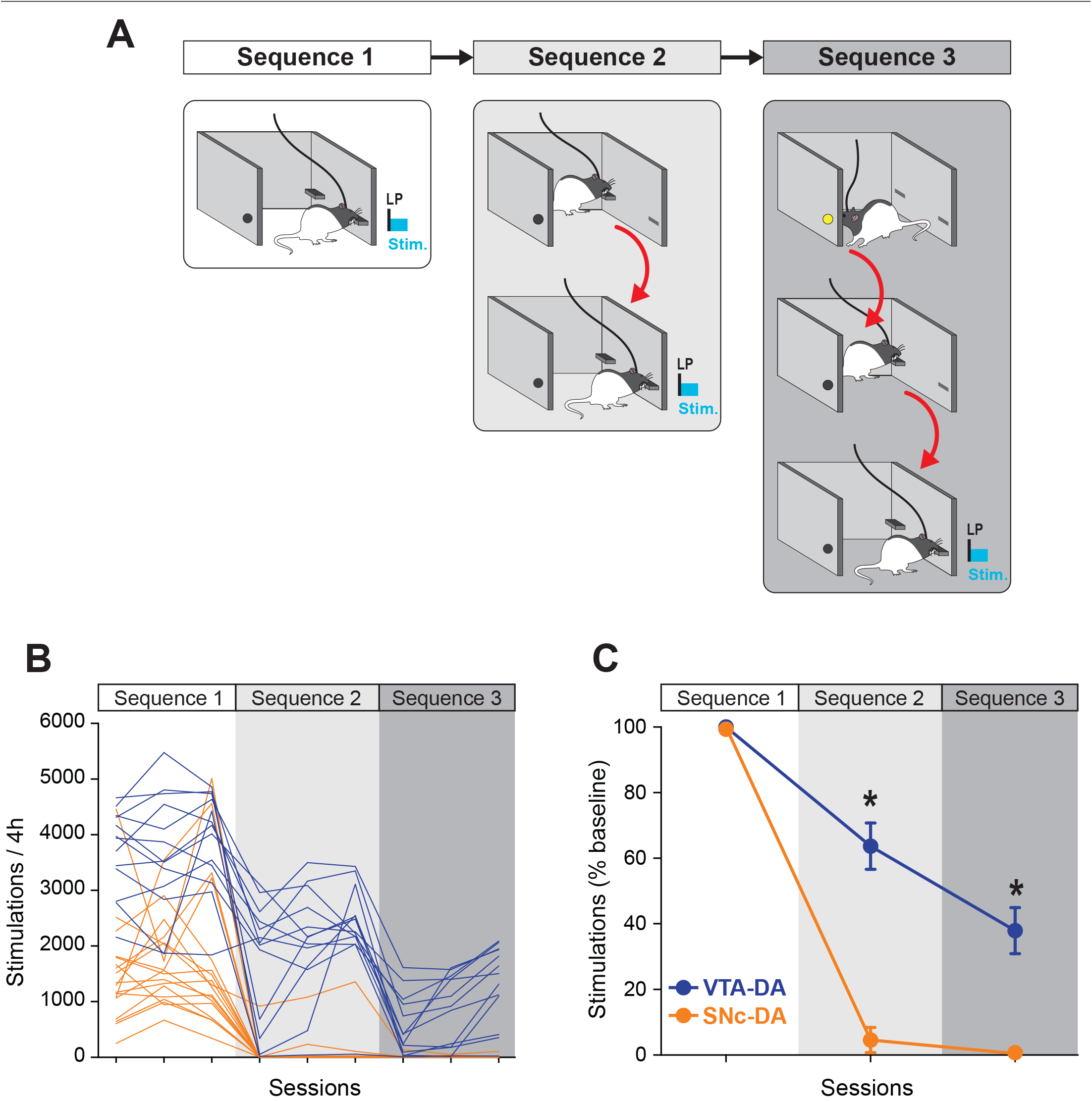
Acquisition of heterogeneous instrumental chain for VTA^DA^, but not SNc^DA^ neuron self-stimulation. **A**. Behavioral paradigm. Rats were initially trained to perform a single instrumental action (press a lever) to obtain an optical stimulation of VTA (n = 14) or SNc (n = 14) dopamine neurons. The instrumental requirement for self-stimulation was then increased to a sequence of 2, then 3, instrumental actions (see text for details). **B**. Number of stimulations obtained under the different instrumental requirements. Each line represents an individual subject. **C**. Average number of stimulations obtained on the final session of each instrumental requirement, expressed as percentage of baseline (baseline = number of stimulation obtained under sequence 1). (*P<0.001, post-hoc Bonferroni t-test). Error bars = SEM

In both groups, the transition from a single action to a two-action sequence induced a sharp decrease in the number of stimulations obtained. However, unlike rats in the SNc group, rats in the VTA group rapidly learned the required sequence and by day 3 obtained ∼65% of the stimulations earned in baseline. Note that some reduction in the number of stimulations is expected even in subjects having perfectly learned the new sequence, as is takes longer to complete a two-action sequence than a single action. Likewise, when the instrumental requirement increased from a two-action to a three-action sequence, the number of stimulations obtained by VTA rats abruptly decreased, but most rats eventually learned the new sequence and by day 3 of this schedule VTA rats obtained ∼40% of their baseline stimulations in the one-lever condition. In contrast, the number of stimulations earned by rats in the SNc group remained extremely low.

To compare the two groups, we conducted a 2-way mixed ANCOVA (Group x Schedule) on the number of stimulations obtained on the last session of each schedule. To account for the difference in baseline performance between VTA and SNc groups, the number of stimulations earned in baseline was used as a covariate. This analysis revealed a main effect of Group (F_1,24_ = 18.006, P<0.001), and Group x Schedule interaction (F_2,48_ = 13.943, P<0.001), confirming the different impact of the increasing action sequence requirement on VTA^DA^ and SNc^DA^ neuron self-stimulation. Thus, VTA dopamine neuron stimulation can bridge the gap between events and guide learning through a series of actions leading to their activation whereas SNc dopamine neurons fail to do so.

## DISCUSSION

The activation of midbrain dopamine neurons, in the VTA or the adjacent SNc, is a potent reinforcer of instrumental behavior^4,5,8,23,24,31^. However, we show here that VTA and SNc dopamine neurons are not functionally equivalent in instrumental reinforcement. Despite superficial similarities in the overt instrumental response, self-stimulation of VTA or SNc dopamine neurons engages different underlying associative structures. Self-stimulation of VTA dopamine neurons was characterized by a high rate of lever pressing, as well as a strong attraction towards the cues/states associated with the stimulation (i.e. the proximity to the active lever, or the active lever itself). These incentive, or “motivational-magnet”, properties acquired by these cues allowed VTA^DA^ self-stimulating animals to easily reengage in the task following a pause (self-imposed or experimentally imposed timeouts) and to flexibly approach the active lever following its spatial relocation. The incentive properties acquired by the stimuli surrounding VTA^DA^ self-stimulation were also evident in the ability of these stimuli to act as conditioned reinforcers, sustaining instrumental responding in the absence of the primary reinforcer (i.e. in absence of optical simulation). This combination of attracting and reinforcing properties exerted by the environmental stimuli surrounding VTA^DA^ stimulation might have facilitated the acquisition of increasingly complex instrumental sequences for the self-stimulation of VTA^DA^ neurons. For instance, the insertion of the “taking” lever simultaneously reinforces the previous action responsible for the apparition of this stimulus (i.e. pressing the “seeking” lever), and attracts the animal towards this new stimulus, the animal ultimately following a gradient of increasing value, or reward-taxis^32^. In contrast, self-stimulation of SNc dopamine neurons was characterized by bouts of high rate of instrumental responding separated by long pauses during which instrumental responding was completely absent. This response pattern suggests that SNc^DA^ stimulation, while capable of reinforcing an instrumental action, fails to confer incentive properties to the cues/states associated with that stimulation. Consistent with this interpretation, rats self-stimulating SNc^DA^ failed to reengage in the task following an imposed timeout, and showed reduced engagement with the active lever following its spatial relocation. Moreover, cues paired with SNc^DA^ stimulation failed to sustain instrumental responding in absence of the stimulation. Consequently, in absence of incentive cues to guide them, rats stimulating SNc^DA^ neurons failed to acquire complex instrumental sequences for SNc^DA^ stimulation.

This study indicates that a limiting factor in SNc^DA^ self-stimulation, and the reason for the overall lower responding in that group, is a reduced motivation to initiate responding. This finding might appear at odds with previous studies (including work from our group) which reported comparable levels of responding for VTA^DA^ and SNc^DA^ stimulation^4,8,23^. Several factors might explain this discrepancy, including i) the nature of the operandum, ii) the novelty of the operandum, and iii) the presence or absence of ancillary stimuli. Indeed, in our previous studies, we used nosepokes as operanda (vs. levers in the present study), which generally allow for higher levels of reinforced, but also spontaneous (non-reinforced) responding^33–35^. Moreover, in those same studies, the nosepoke operanda were made available after animals experienced several Pavlovian conditioning sessions in the same chambers. The introduction of this novel operandum in a safe and familiar environment is likely to increase spontaneous exploration and interaction^36^. Finally, unlike the present study, other studies used ancillary stimuli (during initial training) in the form of brief visual stimuli accompanying VTA^DA^ or SNc^DA^ neuron stimulation^8^. Such response-contingent sensory stimulation is known to maintain non-negligible level of instrumental responding, even in absence of any other primary reward^37,38^. In and of themselves, these factors (type of operandum, novelty of the operandum, and ancillary cues) cannot explain the high rate of responding in VTA^DA^ or SNc^DA^ ICSS experiments, which is clearly due to the reinforcing properties of DA stimulation. However, by ensuring sporadic spontaneous interactions with the instrumental operandum, these experimental conditions might have compensated for the reduced motivation to initiate responding in SNc^DA^ rats and masked potential differences between VTA^DA^ and SNc^DA^ self-stimulation.

Inthisstudy,weinterrogatedVTA^DA^ andSNc^DA^ separately. However, these neural populations do not necessarily function independently from each other. Indeed, VTA^DA^ neurons can potentially influence SNc^DA^ neurons via ascending striato-nigro-striatal loops^39–41^. Therefore, in this study the direct, targeted, optogenetic stimulation of VTA^DA^ neurons might have resulted in an indirect, propagated activation of the SNc^DA^, due to the intrinsic connectivity between these regions but see ^42^. The extent to which the instrumental behavior observed during VTA^DA^ self-stimulation relies strictly on VTA^DA^ activation or reflects the additional recruitment of SNc^DA^ neurons remains to be determined.

Of note, the propagation of the (dopamine) reinforcing signal is a central feature of the TD actor-critic reinforcement model and its proposed neurobiological implementation^43–45^. In this model, the activation of SNc^DA^ neurons and the resulting dopamine release in the dorsolateral striatum (DLS) contributes to (instrumental) response policy learning (the ‘actor’). Conversely, activation of VTA^DA^ neurons and the resulting dopamine release in the ventromedial striatum (VMS) contributes to Pavlovian state-value learning (the ‘critic’). In return, these valued states, encoded in the VMS, can themselves elicit dopamine reinforcement signals, via striato-nigro-striatal loops. Closed loops (VMS → VTA → VMS) presumably allow for the temporal backpropagation of the dopamine reinforcement signal within the ‘critic’ module, allowing animals to identify (or backtrack) the earliest state (or cue) predictive of reward^46^. Open ascending loops (VMS → SNc → DLS) presumably allow for the propagation of the dopamine reinforcement signal from the ‘critic’ to the ‘actor’ module, thereby reinforcing actions that lead to valued states (i.e. actions that increase the prospect of reward). Note the absence of descending loops (DLS → VTA → VMS) in this model.

The potential for the learning-mediated temporal backpropagation of the dopamine reinforcement signal following VTA^DA^, but not SNc^DA^ activation might contribute to the observed behavioral difference between VTA^DA^ and SNc^DA^ self-stimulation. Indeed, the stimulation of VTA^DA^ and the resulting backpropagation of dopamine reinforcement signal might allow for the temporally- and spatially-organized reward seeking behavior observed during VTA^DA^ self-stimulation. In contrast, in absence of backpropagated dopamine signal, the reinforcing effect of SNc^DA^ would be limited to the elemental action that immediately precedes the stimulation (**Fig. S3**). Rats self-stimulating SNc^DA^ neurons might therefore find themselves in a most peculiar situation, avidly engaging in instrumental responding when they, by chance, find themselves in proximity of the active lever, but otherwise showing little motivation to approach the lever or engage in the task.

In conclusion, consistent with prior studies we showed here that the activation of either VTA or SNc dopamine neurons is a potent reinforcer of instrumental behavior^4,8,23,24^. Critically, however, we demonstrate that VTA^DA^ and SNc^DA^ neuron activation produce different “dimensions” of reinforcement, as reflected by the profound behavioral differences observed during VTA^DA^ and SNc^DA^ self stimulation. Indeed, rats self-stimulating VTA^DA^ neurons demonstrated a flexible, organized, and motivated reward-seeking (i.e. stimulation-seeking) behavior. In contrast, rats self stimulating SNc^DA^ neurons, while capable of high rate of stereotyped instrumental behavior during response bouts, appear to lack flexibility and motivation to initiate responding. This demonstrates that the functional specialization of VTA and SNc dopamine neurons is not limited to Pavlovian learning but extends to the instrumental domain. Finally, these results highlight how these parallel, yet interacting, dopamine pathways might contribute to different levels of integration of operant behavior, from hierarchically organized action plans to elemental motor commands^47–50^.

## METHODS

### Animals and Surgeries

Males and females transgenic rats expressing Cre recombinase under control of the tyrosine hydroxylase promoter (Th::cre rats) were used in these studies. Rats were individually housed under a 12 h light/12 h dark cycle, and had unlimited access to food and water except during testing. The majority of the experiments were conducted during the light cycle. All experimental procedures were conducted in accordance with JHU Institutional Animal Care and Use Committees and the US National Institute of Health guidelines. Males and females were distributed as evenly as possible across groups. No significant effects of sex were found; therefore data for males and females were collapsed. Rats (> 300 g males; > 225 g females) were anesthetized with isoflurane (induction 5%, maintenance 1-2%) and received unilateral infusions of AAV5-EF1a-DIO-ChR2-eYFP (titer ∼10^12^) into the VTA or the SNc under stereotaxic guidance. VTA was targeted at AP: -5.4 and -6.2mm from bregma; ML: ± 0.7 from midline; DV: -8.5 and -7.5 from skull). SNc was targeted at AP: -5.0 and -5.8; ML: ± 2.4; DV: -8.0 and -7.0.

This resulted in 4 injection sites for each rat. At each injection site, 0.5 ul of virus was delivered at the rate of 0.1ul/min. Injectors were left in place for 10 min following the infusion to allow for diffusion. Immediately following viral infusions, optic fibers (300um core diameter, 0.37 NA) aimed at VTA (AP: -5.8; ML: ± 0.7; DV: -7.5) or SNc (AP: -5.4; ML: ± 2.4; DV: -7.2) were implanted. Behavioral training started 3-4 weeks post-surgery to allow for gene expression

### Optical Activation

Rats were tethered to optical patch cords (200um) connected to a 473nm blue laser diode (Opto Engine) through a rotary joint (Doric Lenses). Fiberoptic implants and patch cords were constructed in the laboratory and were equipped with a custom lock-in mechanism that ensured secure tethering during long behavioral sessions. An individual stimulation event consisted of a 2s train of light pulses (20 Hz, 40 pulses, 5 ms pulse duration). Unless specified otherwise, the laser output during optogenetic stimulation was 24mW, resulting in an irradiance of ∼8.5 mW/mm^2^/s at the tip of the intracranial fiber (corrected for duty cycle). Light power was verified before and after every behavioral session.

### Intracranial self-stimulation (ICSS)

ICSS sessions were conducted in 12 identical sound-attenuated operant chambers (Med Associates, St. Albans, VT). Operant chambers were fitted with two retractable levers on the front panel, and one nosepoke operandum on the back panel (obstructed in all experiments with the exception of the heterogeneous instrumental chain experiment). During ICSS sessions, a response at the active lever (position counterbalanced) resulted in delivery of a 2 s train of light pulses. Inactive lever responses, and active lever responses occurring during the 2 s light train, were recorded but had no consequence. Ventilation fans provided background noise of 65 dB and a red houselight provided diffuse background illumination during the ICSS sessions.

### Initial acquisition

Before being assigned to the different experiments, all rats were initially trained to acquire instrumental ICSS responding. A minimum of 100 active lever presses per hour, for three consecutive sessions, constituted the criterion for successful acquisition (range: 3-11 sessions). The experiments described here were conducted in four different replication cohorts. The initial acquisition of ICSS was conducted under experimental conditions that differed slightly between cohorts and as a function of the different experiments (sessions’ time in the day/light cycle, session’s duration [1-6h], and presence or absence of an inactive lever), therefore acquisition data were not analyzed. Following initial acquisition, all subsequent experiments were conducted in identical conditions for all cohorts. These differences in procedures during initial ICSS acquisition had no consequences on later behavioral outcomes (no effect of initial training protocol).

### Experiment 1: ICSS response patterns

Rats (VTA n = 20; SNc n = 20) were given 3-5 daily 4-h sessions of ICSS. Response patterns were analyzed for the last session completed.

### Experiment 2: Forced timeout

To determine the influence of forced timeouts on instrumental responding for VTA^DA^ of SNc^DA^ neurons stimulation, a subset of rats included in experiment 1 (VTA n = 15; SNc n =16) went on to complete the following experiment. During a single daily 2-h ICSS session, each response on the active lever resulted in an optical stimulation (2s) and a retraction of both levers (active and inactive) for a period of 12 seconds. At the end of this delay both levers were extended and remained extended until the subsequent stimulation.

### Experiment 3: Lever relocation

Rats (VTA n = 10, SNc n = 8) were trained to lever press for optical stimulation of dopamine neurons in presence of a single (active) lever, in daily 2h sessions. To facilitate the detection of the lever, a discrete cue light (28V) located immediately above the lever remained on during all sessions. After 5-6 training sessions, all rats were tested in 2 probe sessions during which no stimulation was delivered. In one probe session, the lever remained in its usual location. In the other probe session, the panel containing the lever was reconfigured to raise the lever (and the light above the lever) by 8cm. Animals were retrained for 3 sessions between the two probe tests (with the lever in its standard position) and the order of testing was counterbalanced within groups.

### Experiment 4: Cued progressive ratio

Rats were trained to self-stimulate VTA^DA^ (n = 6) or SNc^DA^ (n = 6) neurons, and each stimulation coincided with the presentation of a brief audiovisual cue (white noise and chamber illumination; 0.4s). To increase the relevance of the cue and promote the association between this cue and the optoactivation of dopamine neurons, the response requirement was gradually increased over the course of 8 days, from FR1 to FR5. We noticed that several rats in the SNc^DA^ group had difficulty maintaining responding under these higher response ratio, therefore the response requirement was brought back to RR1.5 or RR2 for the remaining training sessions. To ensure equal number of cue-stimulation pairings in both groups, we imposed a maximal number of stimulations per session, which all animals completed (max. number of stimulation = 200, with the exception of FR3 and FR5 sessions for which the max. number of stimulation was reduced to 100 and 20 respectively). The conditioned incentive properties acquired by the audiovisual cue were then assessed in four progressive ratio (PR) probe tests. Under PR schedule, the number of responses required to earn a stimulation was increased after each stimulation according to the following formula^51^:

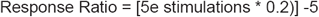

This produced the following response requirement schedule: 1-2-4-6-9-12-15-20-25-32-40-50-62. In two PR test sessions (session 1 and 3), the audiovisual cue continued to be presented on every stimulation (as in training). In the other two PR test sessions (session 2 and 4), the audiovisual cue was contingent on every response on the active lever (responses produced during cue presentation did not prolong the cue). PR probe sessions were separated by 2 days of retraining under the RR1.5 or RR2 schedule described above (**Fig. S2**).

### Experiment 5: Heterogeneous instrumental chain

Rats were initially trained, in 4-h sessions, to press an active “taking” lever to obtain an optoactivation of either VTA^DA^ (n=13) or SNc^DA^ (n=14) neurons. A second, inactive, lever was present at this stage, but had no programmed consequence. After five or six sessions under this reinforcement schedule, rats were required to perform a sequence of two instrumental actions to obtain an optogenetic stimulation. Specifically, rats were required the press a “seeking” lever (corresponding to the previously inactive lever) to gain access to the “taking” lever. A response of the “seeking” caused the insertion of the “taking” lever. A press on the “taking” would then produce the optogenetic stimulation of VTA^DA^ or SNc^DA^ neurons, and the retraction of the “taking” lever. After three sessions under this reinforcement schedule, rats were required to perform a sequence of three instrumental actions to obtain an optostimulation. Specifically, rats were required to perform a nosepoke (on the back panel, opposite to the levers), then press the “seeking” lever, and finally press the “taking” lever to obtain an optostimulation. An LED light located inside the nosepoke operandum signaled that this operandum was available for responding. A nosepoke response caused the termination of this LED light and the insertion of the “seeking” lever. A response of the “seeking” lever caused the insertion of the “taking” lever. A press on the “taking” lever would then produce the optostimulation, as well as the retraction of all levers and the illumination of the nosepoke LED light. Rat were trained on this schedule for three sessions.

Some rats completed more than one experiment. See **Table 1** (Supplemental) for rats’ allocation to the different experiments.

**TABLE 1:**
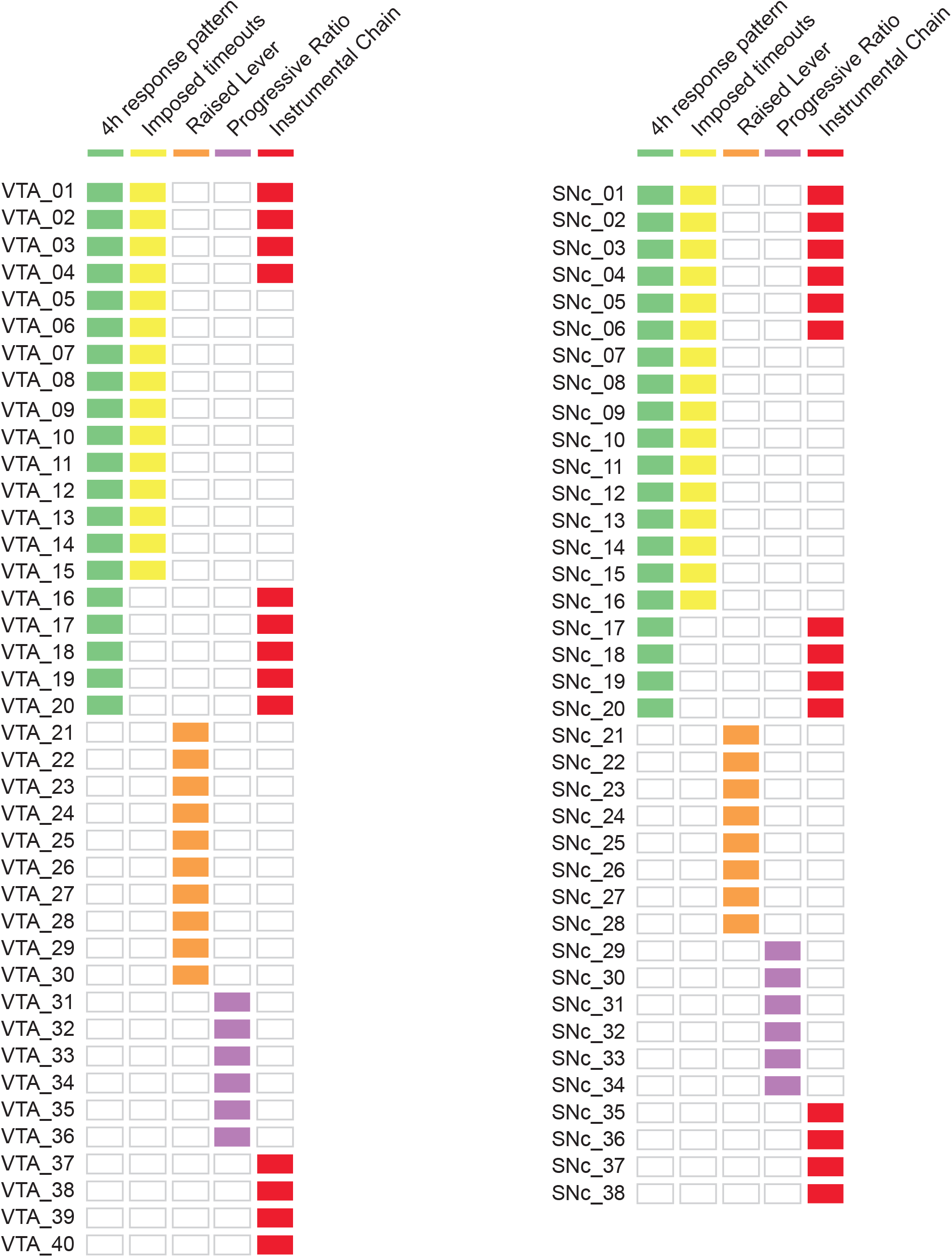
Subjects’ allocation to the different experiments

### Histology

Animals were anesthetized with pentobarbital and perfused intracardially with 0.9% saline followed by 4% paraformaldehyde. Brains were extracted, post-fixed in 4% paraformaldehyde for 24h, and cryoprotected in 30% sucrose for > 48 hours. Brains were then frozen on dry ice and sectioned at 40 mm on a cryostat. Coronal slices were collected onto glass slides and coverslipped with Vectashield mounting medium with DAPI. Fiber tip position and eYFP-ChR2 virus expression were verified under a fluorescence microscope (Zeiss Microscopy, Thornwood, NY).

### Statistical Analysis

Statistical analyses were performed using the SPSS statistical package (IBM SPSS Statistics, Version 25.0.0.1). Graphs were generated using GraphPad Prism 9 and MATLAB R2016a (MathWorks). For each data set, Lilliefors and Brown-Forsythe tests were run to test for normality and equal variance, respectively. When appropriate, parametric tests were conducted and consisted generally of mixed-design repeated-measures RM ANOVAs with Group (VTA^DA^ or SNc^DA^) as between-subject factor and Session as within-subject factor. Post-hoc and planned comparisons were carried with two-tailed Student’s t-tests. When non-normality and/or unequal variances were observed in our data set, nonparametric tests were conducted and consisted of Mann-Whitney U test, or Wilcoxon signed-rank Z test (for between-or within-subjects comparison respectively). Significance was assessed against a type I error rate of 0.05. Additionally, for each pairwise or independent comparison, we provide a 95% confidence interval of the difference of means, which was determined by a bias-corrected and accelerated (BCa) bootstrapped method, (free from distributional assumptions) with 5000 resamples. Effect size were estimated with the rank-biserial correlation.

## Supplemental Figures

**FIGURE S1:**
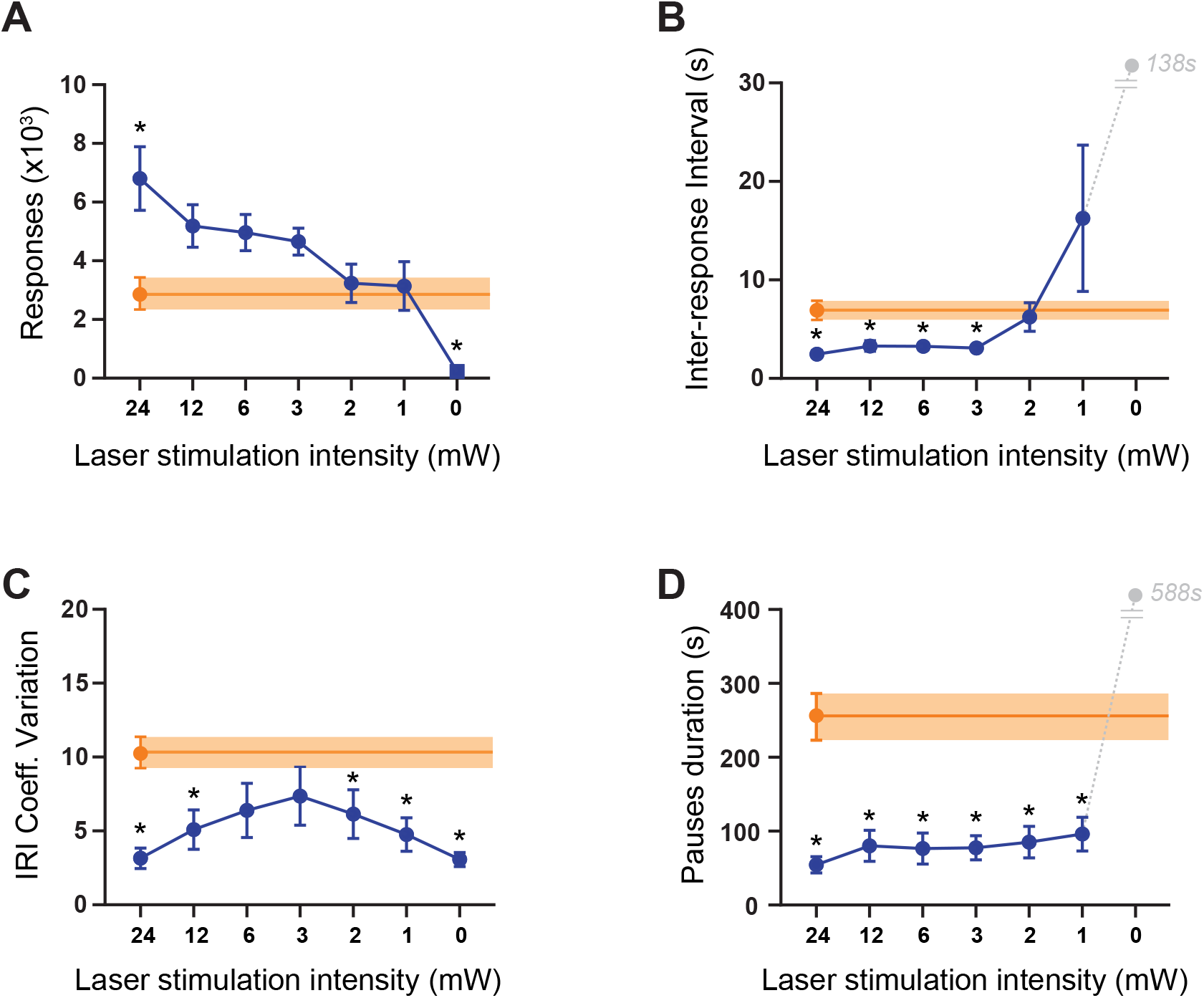
Operant responding for different intensities of VTA^DA^ neuron stimulation. **A**. Total responses per session, **B**. Inter-response interval (IRI), **C**. Coefficient of variation of IRIs within a session, **D**. Average pause duration. Reducing VTA^DA^ stimulation intensity reduced the number of responses and increased the average IRI, but those lower stimulation intensities failed to reproduce the irregular pattern or long pauses in responding that characterize SNc^DA^ self-stimulation. Error bars and error bands = SEM. Orange line and shading represent the average ± SEM for the SNc group at the maximum intensity of 24mW. (*denote significant difference from SNc^DA^ 24W, P<0.05 bootstrapped t-tests).

**FIGURE S2:**
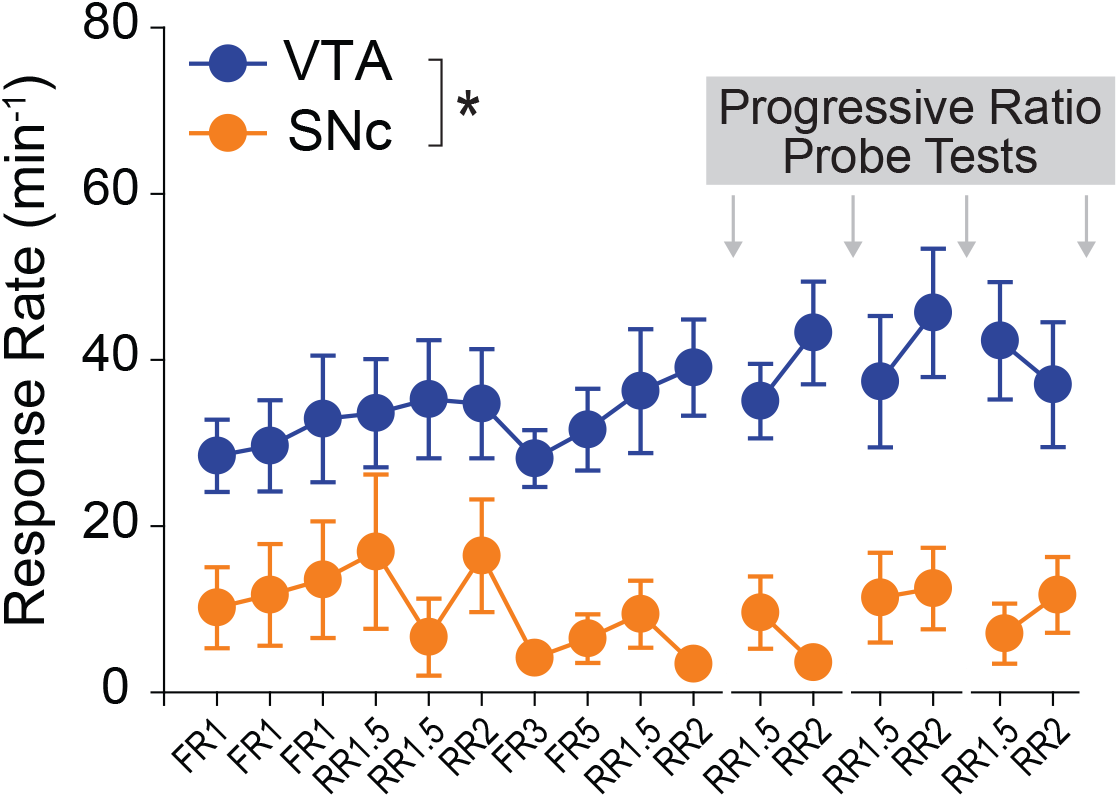
Rate of responding for VTA^DA^ or SNc^DA^ stimulation under different reinforcement ratios. Rats initially acquired instrumental self-stimulation of VTA^DA^ or SNc^DA^ neurons under a continuous schedule of reinforcement (FR1). A brief audio-visual cue accompanied each optostimulation. In an attempt to strengthen the association between this cue and the optostimulation, the response requirement was progressively increased (FR1, RR2, FR3, and FR5); thereby reducing the response-stimulation contingency, while maintaining a maximal cue-stimulation contingency. Because several rats in the SNc^DA^ group had difficulty maintaining responding under these higher response ratio, the response requirement was brought back to RR1.5 or RR2 for the remaining training sessions. All rats obtained the maximum number of stimulation on each session, although the VTA^DA^ rats had a higher response rate (*Main Group effect F1,10 = 13.86, P=0.004).

**FIGURE S3:**
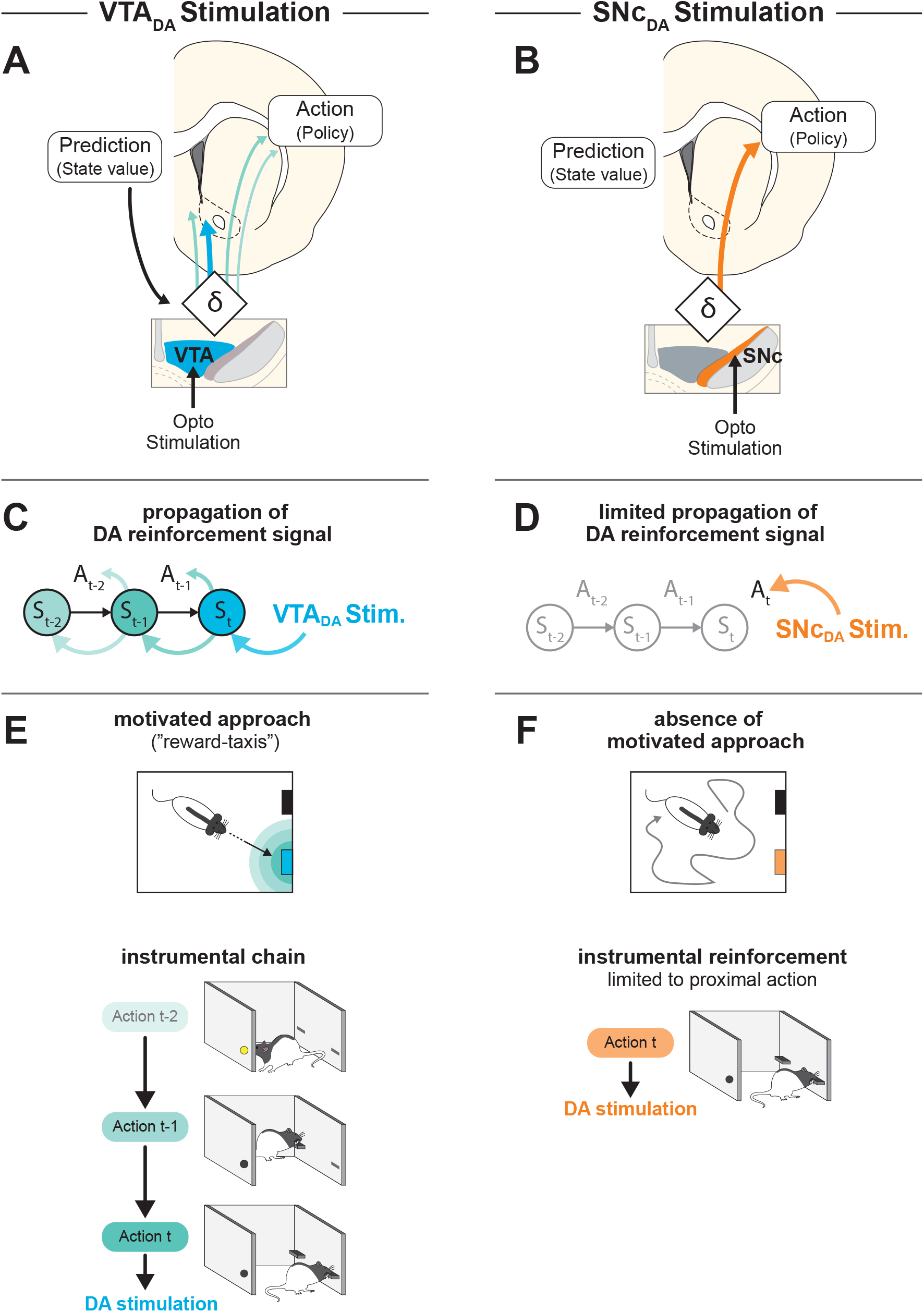
Proposed mechanism for VTA^DA^- and SNc^DA^-mediated reinforcement. **A-B**. Actor-Critic circuit motif. State values are proposed to be encoded in the ventromedial striatum (VMS, the critic). Policies, or action values are proposed to be encoded in the dorsolateral striatum (DLS, the actor). VTA and SNc dopamine neurons provide reinforcement signals to VMS and DLS respectively. Feedback projections from the ventromedial striatum allows valued states to activate VTA and SNc dopamine neurons resulting in the propagation of the dopamine reinforcement signal. For simplicity, the midbrain microcircuitry is not shown. **C-D**. This circuit motif predicts that stimulation of VTA^DA^ produces a direct reinforcement signal to the critic (blue arrow), but also temporally and anatomically propagated reinforcement signals to both the actor and the critic modules (green arrows). In contrast, stimulation of the SNc^DA^ produces a direct reinforcement signal to the actor (orange arrow) with little potential for backpropagation of that reinforcement signal. **E-F**. Consequences for behavior. See discussion for details

